# Regulation of XPG-DNA damage binding dynamics by pre- and post-incision nucleotide excision repair factors and EXO1

**DOI:** 10.1101/2024.12.06.627128

**Authors:** Alba Muniesa-Vargas, Cristina Ribeiro-Silva, Bart Geverts, Carlota Davó-Martínez, Jacinta van de Grint, Maroussia Ganpat, Karen L. Thijssen, Joris Pothof, Adriaan B. Houtsmuller, Arjan F. Theil, Wim Vermeulen, Hannes Lans

## Abstract

The XPG endonuclease plays a crucial role in nucleotide excision repair (NER) and other genome maintenance pathways. Precise regulation of XPG recruitment and activity during DNA repair is essential to avoid erroneous DNA incisions and genomic instability. In this study, we employed live-cell imaging to investigate how XPG function is regulated during NER, focusing on its dynamic interactions with key factors involved in the pre- and post-incision steps. We found that TFIIH and XPA facilitate the recruitment and association of XPG with DNA damage, and that XPG localizes separately from TFIIH to UV-induced lesions. Furthermore, our results show that XPG’s dissociation from DNA damage is triggered by its own incision activity as well as by that of XPF. Additionally, the exonuclease EXO1 promotes XPG dissociation, likely by processing incised DNA, even in the absence of XPG-mediated incision. Our findings help to better understand the regulatory mechanisms that control XPG activity during NER and provide important insights into the complex dynamics of the repair process.

## Introduction

DNA damage is a constant threat to cells. It is estimated that cells incur thousands of new DNA lesions every day from external sources, such as UV light from the sun, and from reactive chemicals released during metabolism (1, 2). Cells rely on the DNA damage response (DDR) to maintain genome integrity. This is an evolutionarily conserved, intricate network of DNA repair, chromatin remodeling, DNA damage signaling, and tolerance mechanisms to cope with DNA lesions (3). Disruption of the DDR and inefficient removal of lesions cause genome instability and have disastrous consequences for organisms, leading to genetic diseases, developmental failure, premature aging, and cancer susceptibility (4).

XPG/ERCC5 is a major DDR endonuclease and an essential factor in maintaining genome integrity. XPG is best known for its role in nucleotide excision repair (NER), but functions in multiple other genome maintenance mechanisms as well (5), such as R-loop resolution (6–8), replication stress and homologous recombination (9, 10), and base excision repair (11–13). XPG harbors two nuclease domains termed N and I, which are separated by a spacer region, two highly conserved domains, i.e., the D1 and D2 boxes, and a PIP box with which it was reported to interact with PCNA (14). Hereditary mutations in XPG cause an extraordinary broad spectrum of clinical features, associated with rare human disorders such as xeroderma pigmentosum (XP) and XP with additional Cockayne syndrome features, called Xeroderma pigmentosum Cockayne syndrome (XPCS) complex or (in the more severe form) cerebro-oculofacio-skeletal (COFS) syndrome (5, 15). XP is characterized by photosensitivity and pigmentation abnormalities of the skin, and high incidence of skin cancer (16). XPCS and COFS patients display mental retardation, developmental delay, progeria and severe, progressive neurological abnormalities. To understand the pathogenesis of these diseases and why different mutations cause different symptoms, it is important to precisely understand how XPG function is controlled in NER and other DDR pathways.

NER is a highly coordinated DNA repair pathway that removes many different types of bulky lesions, such as the UV light-induced cyclobutane-pyrimidine dimers (CPDs) and 6-4 pyrimidine-pyrimidone photoproducts, chemotherapy-induced intrastrand crosslinks and ROS-induced cyclopurines (17). This versatility relies on its ability to recognize similar helix distortions in DNA inflicted by different types of DNA damage, instead of the lesions themselves. NER detects and removes DNA damage by either one of two subpathways: global genome-NER (GG-NER) and transcription-coupled NER (TC-NER). GG-NER is initiated by damage detection anywhere in the genome by the coordinated action of DDB2, which is part of the E3 ubiquitin ligase complex CRL4^DDB2^, and XPC, which is part of the trimeric XPC-RAD23B-CETN2 complex (18). TC-NER is initiated when damage in the template strand blocks forward movement of RNA polymerase II during transcription elongation (19). Stalling of RNA Polymerase II at DNA damage leads to stable binding of the translocase CSB and subsequent recruitment of the E3 ubiquitin ligase CRL4^CSA^ and the factors STK19 and UVSSA/USP7, involving the RNA Polymerase II-bound ELOF1 protein as well.

Damage detection by either subpathway leads to the sequential and cooperative buildup of the core NER incision complex, which is precisely coordinated by multiple protein-protein interactions and posttranslational modifications (20). In GG-NER, the 10-subunit complex TFIIH is recruited through interaction of its XPB and p62 subunits with XPC bound to DNA lesions (21, 22). This in turn coincides with DDB2 dissociation and promotes the transition from temporary to stable binding of XPC to DNA damage (23). In TC-NER, TFIIH is recruited and positioned by interactions with UVSSA (24) and STK19 (25–28). TFIIH unwinds and scans the DNA with its helicase XPD and translocase XPB to verify the presence of lesions (20). This TFIIH activity is stimulated by the DNA binding protein XPA (29–34). Thus, TFIIH recruitment precedes that of XPA (35–37), but XPA in turn stabilizes TFIIH association with damaged DNA (23). Also, RPA is recruited, which binds to the ssDNA opposite of the damaged strand and interacts with XPA to stabilize its DNA damage binding (38–40). The helicase and translocase activities of TFIIH generate a DNA bubble necessary for the ss/dsDNA junction binding and incision activity of endonucleases ERCC1-XPF and XPG. These incise DNA, respectively, 5’ and 3’ of the lesion, leading to removal of the damaged DNA strand. ERCC1-XPF is recruited via an interaction of ERCC1 with XPA (41–43), and will only incise DNA in the presence of XPG (44). While both ERCC1-XPF and XPG are thought to be positioned through an interaction with RPA (45), it is still not entirely clear how XPG recruitment is regulated. Previous immunoprecipitation experiments indicated that XPG can form a stable complex with TFIIH that is able to incise DNA bubble substrates *in vitro* (46), suggesting that XPG already accompanies TFIIH when this complex binds to DNA damage. This model, however, contrasts cellular mobility studies that suggested that XPG diffuses freely through the nucleus without being part of a larger complex (47).

Additionally, it is unclear when and how XPG dissociates from DNA damage. Following DNA incision, the HLTF translocase promotes the dissociation of the damaged DNA strand and of several incision complex factors, including XPG (48). Excision assays showed a tight association of XPG and TFIIH with the excised DNA product, suggesting that XPG dissociates simultaneously with TFIIH upon incision (49). In apparent contradiction, *in vitro* reconstitution of dual incision of cisplatin-DNA adducts suggested that after DNA incision XPG first promotes the recruitment of downstream DNA replication factors RFC1 and PCNA, together with RPA, to initiate DNA synthesis (50) and restore the resulting 22-30 nucleotides DNA gap. Interestingly, this DNA repair synthesis appears to be cell cycle-dependent, as loading of DNA polymerases δ and κ by PCNA is favored in non-S phase cells, while loading of polymerase ε only happens in S phase cells (51, 52). Moreover, the 5’-3’ exonuclease EXO1 has been suggested to compete with RFC1 and PCNA-mediated DNA repair synthesis, by resecting DNA and generating long ssDNA structures that activate the DNA damage checkpoint (53). Also, this was reported to be dependent on the cell cycle phase, since recruitment of a fluorescently-tagged EXO1b isoform to UV damage was only observed in non-S phase cells (54). It is still ambiguous if and how XPG dissociation is regulated by these post-incision factors and whether the cell cycle plays a role in the crosstalk and coordination between XPG and the post-incision step.

To better understand the dynamic activity of XPG during NER, and its function and interplay with other NER factors, here we investigated, by live-cell imaging of endogenous XPG in human cells, how XPG association to and dissociation from DNA damage is organized within the NER incision complex and after incision has taken place. We show that while XPG recruitment depends on TFIIH, XPG is separately recruited to sites of UV damage. Furthermore, XPG dissociation is stimulated by the catalytic activity of XPF, of XPG itself and by EXO1 activity, irrespective of whether XPG itself can incise DNA or not.

## Materials and methods

### Cell lines, culture conditions and treatments

All human osteosarcoma U2OS cells generated and used are listed in Supplementary Table 1. Cells were cultured at 37°C in a humidified atmosphere and 5% CO_2_ in a 1:1 mixture of DMEM (Lonza) and Ham’s F10 (Lonza) supplemented with 10% fetal calf serum (FCS) and 1% penicillin-streptomycin (PS). All cell lines were regularly tested for mycoplasma contamination. To knock-in mAID-mClover at the C-terminus of *XPG*, cells were transfected with pLentiCRISPR-v2 (55) carrying an sgRNA (5’-AAGGAAACTAAGACGTGCGA-3’) and two plasmids carrying mAID-mClover flanked by XPG sequences and a neomycin casette (pMK289, Addgene plasmid #72827) or a hygromycin cassette (pMK290, Addgene plasmid #72828). To knock-in mAID-mClover at the C-terminus of *XPD*, cells were transfected with Cas9 protein, a synthetic guide RNA targeting XPD (5’-GGCAAGACTCAGGAGTCACC-3’) and a plasmid carrying mAID-mClover flanked by XPD sequences. To knock-in FLAG-mScarlet at the N-terminus of *PCNA*, cells were transfected with pLentiCRISPR-v2 carrying an sgRNA (5’-CCACTCCGCCACCATGTTCG-3’) and a plasmid expressing FLAG-mScarlet flanked by PCNA sequences. To knock-in TEV-mAID-mClover at the C-terminus of *XPF*, cells were transfected an sgRNA targeting XPF (5’-CCGCTGAAAAGTACAGGCAT-3’) and a plasmid expressing mClover flanked by XPF sequences. *XPC* was knocked out by transient transfection of cells with pLentiCRISPR-V2 plasmids carrying an sgRNA (5’-GCTCGGAAACGCGCGGCCGG-3’) targeting exon 1 and a plasmid carrying a blasticidin selection cassette flanked by XPC sequences, which upon knock-in disrupts XPC expression. XPC KO cells that had this construct knocked-in at the N-terminus of XPC were isolated using blasticidin selection. *XPA* was knocked out by transient transfection of cells with pLentiCRISPR-V2 plasmids expressing the sgRNAs 5’-GGCGGCTTTAGAGCAACCCG-3’ and 5’-GTATCGAGCGGAAGCGGCAG-3’, and carrying a puromycin cassette. XPA KO cells were isolated using puromycin selection. To generate E791A catalytically dead XPG mutant cells, cells were transfected with the Cas9 protein, a synthetic guide RNA targeting XPG (5’-CCAGGCTCCCATGGAAGCAG-3’), and a DNA oligomer of 59 bases used as a repair template containing the mutation 5’-TCCCTACATCCAGGCTCCCATGGAAGCAGCCGCGCAGTGCGCCATCCTGGACCTGACTG-3’). Transfected cells were grown in clones and correct generation of the mutation was confirmed using PCR and restriction with ApeKI. ApeKI restriction site was introduced changing AG > CC leading to the E791A mutation, followed by sequencing. All sgRNAs and Cas9 protein were obtained from Integrated DNA Technologies. All transfections with sgRNA were done using Lipofectamine CRISPRMAX Cas9 (Invitrogen) and plasmid transfections were performed using JetPEI (Promega) according to manufacturer’s instructions.

### siRNA treatments

Cells were transfected with siRNAs 48 h prior to each experiment using RNAiMax (Invitrogen), according to the manufacturer’s instructions. siRNA efficiency was validated by immunoblotting (Supplementary Figure 1A, B, and Supplementary Figure 2C, D, E and G). siRNA oligomers were purchased from Thermo Scientific Dharmacon: CTRL 5’UGGUUUACAUGUCGACUAA-3’ (D-001210-05-20), p62/GTF2H1 (L-010924-00, Smart Pool), XPA 5’-CUGAUGAUAAACACAAGCUUA-3’ (MJAWM-000011); XPF 5’-AAGACGAGCUCACGAGUAU-3’ (D-019946-04); PCNA 5’-GUGUAAACCUGCAGAGCAU-3’; EXO1 5’-GCACGUAAUUCAAGUGAUG-3’; RFC1 (M-009290-01, Smart Pool)

### Colony forming assays

Cells were seeded in triplicate in 6-well plates. We seeded 400 cells/well for U2OS DIVa and U2OS DIVa XPG-mClover, and 600 cells/well for U2OS DIVa XPG(E791A)-mClover. Cells were treated with the indicated doses of UV-C 24 h after seeding. After 7-10 days, colonies were fixed and stained with 1 % w/v Coomassie Blue (Bio-Rad) dispersed in a 50% Methanol, 10% Acetic Acid solution. Colonies were counted with the integrated colony counter GelCount (Oxford Optronix).

### UV-C irradiation

For global UV-irradiation, cells were grown on coverslips, washed with PBS, and irradiated with UV-C using a germicidal lamp (254 nm; TUV lamp, Phillips) at the indicated doses. For iFRAP experiments, local UV-damage (LUD) was induced by applying 80 J/m^2^ of UV-C irradiation through a 5 μm polycarbonate filter (Millipore) that was placed on top of a monolayer of cells. For LUD in immunofluorescence experiments, 60 J/m^2^ of UV-C irradiation through an 8 μm polycarbonate filter was applied.

### Fluorescence Recovery After Photobleaching (FRAP)

Cells were grown on coverslips and imaged with a Leica TCS SP5 microscope using a 40x/1.25 NA HCX PL APO CS oil immersion lens (Leica microsystems) at 37°C and 5% CO_2_. FRAP was performed as previously described (56). In short, the nuclear fluorescence signal of mClover-tagged proteins was measured in a strip across the nucleus (512×16 pixels, zoom 12 x) at 400 Hz of a 488 nm laser until a steady fluorescence level was measured (pre-bleach). The fluorescence signal in the strip was then bleached with high laser power and the recovery of fluorescence was subsequently monitored. Fluorescence signal was normalized to the average pre-bleach fluorescence after background subtraction. The average immobile fractions (F_imm_) in the FRAP curves were calculated using the fluorescence intensity immediately after UV (I_0_) and the average final fluorescence signal over the last 20 s of the FRAP measurements, from untreated (I_final, unt_) and UV-treated cells (I_final, UV_), applying the formula:

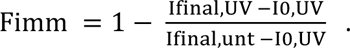

### Monte Carlo simulations

Experimental FRAP curves were fit to computer simulation-generated FRAP curves, as previously described (57, 58). Simulated FRAP curves were computed based on a model that simulates the diffusion of molecules, their binding to and dissociation from immobile elements (i.e. DNA), in an ellipsoidal volume (i.e. the nucleus).

### Inverse Fluorescence Recovery After Photobleaching (iFRAP)

iFRAP was performed as described (59) on a Leica SP8 microscope using a 40/1.25 NA HCX PL APO CS oil immersion lens (Leica microsystems) at 37°C and 5% CO_2._ Cells were grown on coverslips and LUD was induced by irradiating cells through a microporous filter. Next, the fluorescence signal of mClover-tagged proteins was monitored over time in three areas in the cell: in the LUD, in a non-damage nuclear area, and in a cytoplasmic area (background), while the rest of the nucleus was bleached with high laser power. Fluorescence levels in the LUD were normalized to the pre-damage conditions after background subtraction (cytoplasmic area). The non-damaged area served as internal control and was not plotted. To obtain the half-time of protein residence in the LUD the non-linear regression fitted to one-phase exponential decay analysis was applied to the iFRAP curves, using Graph Pad Prism version 9 for Windows (GraphPad Software, La Jolla California USA).

### UV-C laser irradiation (for real-time accumulation)

To study real-time accumulation kinetics of mClover-tagged proteins to laser—induced local UV-C irradiation in living cells, a 2 mW pulsed (7.8 kHz) diode-pumped-solid state laser emitting at 266 nm (Rapp Optoelectronic, Hamburg GmbH) coupled to a Leica SP8 confocal microscope was used. Cells were grown on quartz coverslips and irradiated through an Ultrafluar quartz 100x/1.35 NA glycerol immersion lens (Carl Zeiss Micro Imaging Inc.). Accumulation curves were normalized to pre-damage fluorescence signal, set at 1.

### Immunoblotting

Cells were washed twice with PBS prior to protein extraction using a 1:1 mixture of PBS and 2x Concentrate Laemmli buffer (Merck Sigma-Aldrich) containing 1:1000 benzonase (Millipore). Protein lysates were boiled at 98°C for 5 min, separated in SDS-PAGE gels and transferred onto PVDF membranes (0.45 µm, Merck Millipore). Membranes were incubated in 5% BSA for 1 h and subsequently incubated with primary and fluorescently-labeled secondary antibodies (Supplementary Tables S2 and S3) for 1-2 h at room temperature, or at 4°C overnight. Protein bands were visualized and analyzed with the Odyssey CLx Infrared Imaging System (LI-COR Biosciences).

### Unscheduled DNA synthesis (UDS)

Cells were seeded on 24 mm coverslips and treated with siRNAs 48 h before UV-C irradiation through an 8 μm polycarbonate filter. Cells were labeled with 10 µM ethynyl-2’-deoxyuridine (EdU, Invitrogen) for 30 min. Cells were fixed in 2% paraformaldehyde and permeabilized in 0.1% Triton in PBS. For CPD staining, DNA was denatured with a 70 mM NaOH solution in PBS for 5 min. To visualize EdU, coverslips were incubated with a Click-IT buffer containing Alexa Fluor 647 azide (60 μΜ, Invitrogen), Tris-HCl (50Mm), CuSO_4_ * 5H_2_O (4 mM Sigma) and ascorbic acid (10 mM Sigma) for 1h.

### Immunofluorescence

Cells were seeded on 24 mm coverslips, labeled with 10 µM EdU for 30 min, when indicated, fixed in 2% paraformaldehyde and permeabilized in 0.1% Triton in PBS. For CPD staining, DNA was denatured with a 70 mM NaOH solution in PBS for 5 min. Blocking was performed with 3% BSA and 2.25% glycine in PBS-T (0.05% Tween) for 1 h followed by 2 h incubation at room temperature with primary antibodies and 1 h with secondary antibodies and DAPI (Sigma) staining. Prior to antibody staining, EdU was visualized by incubating the coverslips with a Click-IT buffer containing Alexa Fluor 647 azide (60 μΜ, Invitrogen), Tris-HCl (50Mm), CuSO_4_ * 5H_2_O (4 mM Sigma) and ascorbic acid (10 mM Sigma) for 1h. Coverslips were mounted on slides using Aqua-Poly/Mount (Polysciences, Inc.). A Zeiss LSM700 microscope equipped with 40x Plan-apochromat 1.3 NA oil immersion lens (Carl Zeiss Micro Imaging Inc.) was used to acquire images. Protein recruitment to UV lesions was quantified by dividing the average fluorescence signal intensity at LUD by the average nuclear fluorescence intensity, as measured using FIJI image analysis software.

### Statistical Analysis

Mean values and S.E.M. error bars are shown for each experiment. For the statistical significance analysis of immobile fractions and immunofluorescence at LUDs, we applied a One-Way ANOVA with correction for multiple comparison. For iFRAP assays a Non-linear regression analysis was performed. All analyses were performed using Graph Pad Prism version 9 for Windows (GraphPad Software, La Jolla California USA).

## Results

### XPG recruitment is promoted by TFIIH and XPA

To study the dynamic interactions of XPG with DNA damage during NER, we used fluorescence microscopy to determine how XPG assembly to and disassembly from the initiation complex is organized in cells. To this end, we inserted the fluorescent protein tag mClover (together with the mAID degron tag, which is not used in this study, thus for simplicity we refer to it as XPG-mClover) at the C-terminus of the *XPG/ERCC5* locus in U2OS DIvA cells (60) using CRISPR/Cas9. Successful XPG-mClover knock-in (KI) was validated by fluorescence microscopy and by immunoblot, showing clear nuclear localization of XPG-mClover and exclusive expression of the fusion protein (Figure 1A, B). Functionality of mClover-tagged XPG was confirmed in a UV colony survival assay, showing that the UV sensitivity of XPG-mClover KI cells was comparable to that of the parental cell line (Figure 1C).

**Figure 1.**
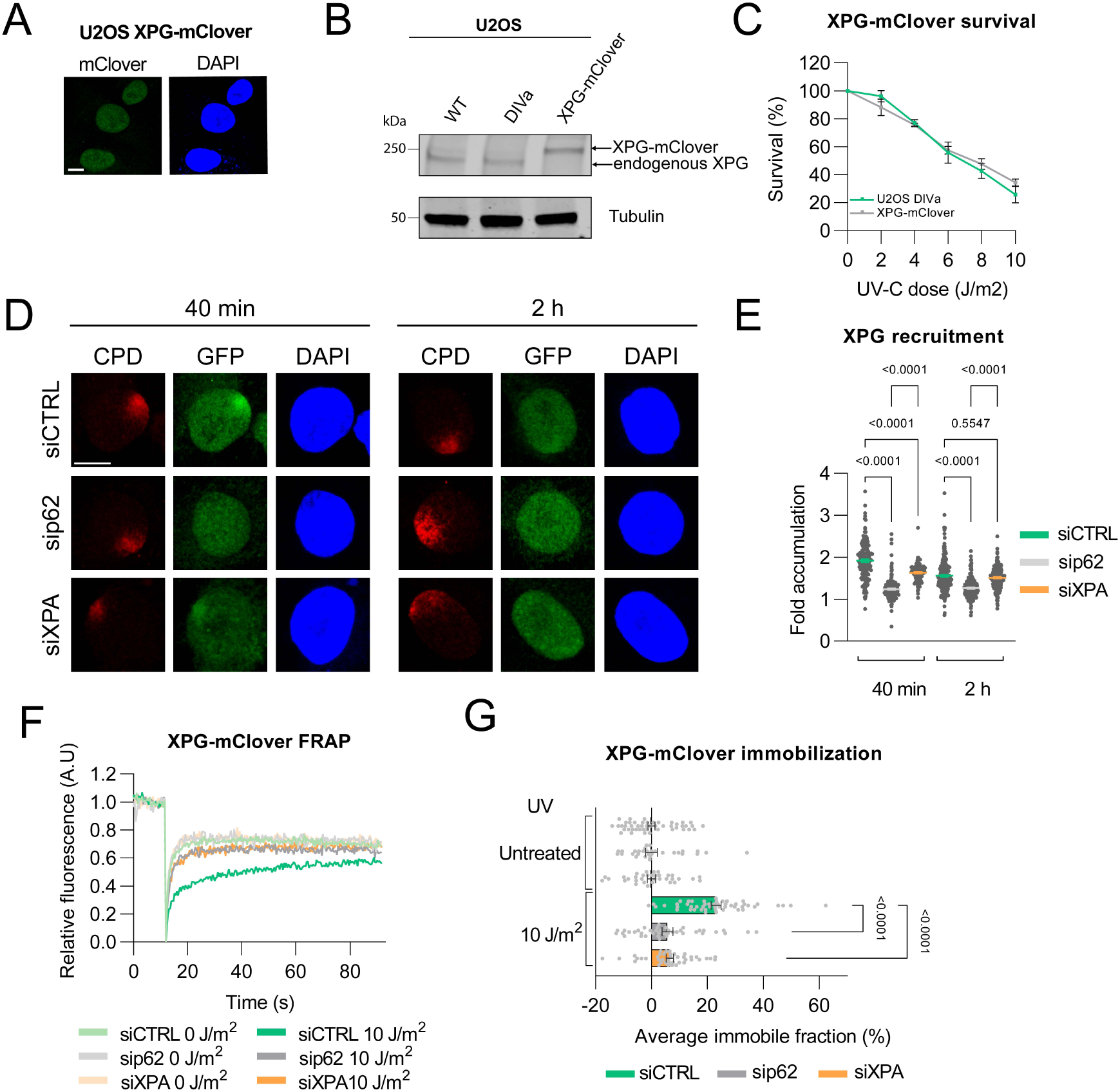
XPG recruitment and binding to DNA damage depends on TFIIH and XPA. (**A**) Representative image of fixed U2OS DIVa XPG-mClover knock-in cells. DNA was stained with DAPI. Scale bar represents 10 µM. (**B**) Immunoblot analysis of wild type (WT), U2OS DIVa and XPG-mClover knock-in U2OS DIVa cells. Immunoblot was stained with antibodies against XPG and tubulin (loading control). (**C**) UV-C colony survival of U2OS DIVa and U2OS DIVa XPG-mClover knock-in cells after UV irradiation at the indicated doses. Survival was plotted as the percentage of colonies obtained after treatment compared to the mean number of colonies in untreated conditions, set as 100%. Mean & SEM of three independent experiments, each performed in triplicate. (**D**) Representative immunofluorescence images of XPG-mClover knock-in U2OS DIVa cells, treated with control, p62 or XPA siRNA, 40 min or 2 h after UV irradiation with 60 J/m^2^ UV-C through a microporous filter to induce LUD. Images show XPG-mClover accumulation at LUD marked by anti-CPD staining. DNA is stained with DAPI. Scale bar 10 µm. (**E**) Quantification of XPG-mClover at LUD as described in (D). The fold accumulation was calculated by normalizing fluorescence intensity at sites of local damage to the nuclear background and plotted as the average of the number of cells per condition from three independent experiments. Number of cells: n= 159, 207, 104 (for 40 min siCTRL, sip62, siXPA); n=246, 248, 228 (for 2 h siCTRL, sip62, siXPA). Numbers in the quantification graphs represent *p-values* determined by ONE-WAY ANOVA. Error bars represent SEM. (**F** and **G**) FRAP analysis of endogenous XPG mobility in U2OS DIVa XPG-mClover cells and quantification of average immobile fraction, over the last 20 s, in cells treated with control (CTRL), p62 and XPA siRNA, non-irradiated (0 J/m^2^) and immediately after UV irradiation (10 J/m^2^). Number of cells: n= 47, 51 (for siCTRL 0 and 10 J/m^2^); n=32, 40 (for sip62 0 and 10 J/m^2^); n=34, 41 (for siXPA 0 and 10 J/m^2^). For all curves, XPG-mClover fluorescence recovery was measured in a strip across the nucleus after bleaching and normalized to the bleach depth. Each FRAP curve represents the average of four independent experiments. Numbers in the quantification graph represent *p-values* determined by ONE-WAY ANOVA. Error bars represent SEM.

To further validate the functionality of mClover-tagged XPG, we tested its recruitment to DNA damage using immunofluorescence after induction of local UV damage (LUD) through a microporous filter. CPD staining was used as a marker of UV damage, as CPDs are slowly repaired (61). 40 min after UV irradiation, we observed clear accumulation of XPG-mClover at UV-induced DNA lesions, which had diminished 2 h after UV due to ongoing repair (Figure 1D, E). Previous similar immunofluorescence studies in patient fibroblasts indicated that stable XPG recruitment depends on functional TFIIH (47, 62). We confirmed this by showing strongly decreased XPG recruitment to DNA damage after siRNA-mediated depletion of the TFIIH subunit p62 (sip62) (Supplementary Figure 1A), 40 min and 2 h after UV irradiation (Figure 1D, E). In addition, *in vitro* dual excision assays suggested that XPA binding to TFIIH and DNA damage is needed for stable assembly of other incision complex factors (63). Indeed, we found that XPG recruitment to DNA damage was significantly reduced, though not fully impaired, in cells depleted of XPA (siXPA) (Figure 1D, E; Supplementary Figure 1B). These results confirm that endogenously tagged XPG is functional and behaves similarly compared to non-tagged XPG.

### XPG localizes separately from TFIIH to DNA damage

To further investigate the dynamic interaction of mClover-tagged XPG with damaged chromatin, we measured its nuclear mobility before and after DNA damage induction using fluorescence recovery after photobleaching (FRAP). In FRAP, incomplete fluorescence recovery reflects immobilization of a fraction of the mClover-tagged molecules, which after UV irradiation is indicative of their stable association to DNA damage (64). In cells treated with control siRNA (siCTRL), a significant fraction of XPG-mClover molecules became immobile immediately after UV irradiation (Figure 1F, G). However, in cells in which p62 or XPA was depleted this UV-induced immobilized fraction was strongly reduced, indicating that both TFIIH and XPA promote binding of XPG to DNA damage. This FRAP reflects the recruitment and dynamic association of XPG with DNA damage immediately after UV irradiation. Combined with the immunofluorescence experiments (Figure 1D, E), which reflect the steady recruitment 40 min after UV irradiation, these results suggest that TFIIH is essential for recruitment of XPG to DNA damage. XPA, on the other hand, appears not essential for recruitment of XPG per se, but rather promotes its association with DNA damage upon recruitment.

XPG was previously reported to form a stable complex with TFIIH (46). We were therefore interested in testing if indeed XPG diffuses together with TFIIH and whether they bind to sites of DNA damage as a complex or separately. To compare the association of XPG with DNA damage to that of TFIIH, we fused the mClover tag (together with the mAID degron tag, which is not used in this study) to the C-terminus of TFIIH subunit XPD in U2OS cells using CRISPR/Cas9 (Supplementary Figure 1C, D) and performed FRAP experiments following UV-irradiation. This showed that a significant fraction of XPD-mClover molecules were bound to DNA damage shortly after UV (Figure 2A), comparable as observed with XPG-mClover (Figure 2B). Because immunofluorescence and FRAP experiments indicated that XPG association to DNA damage is stimulated by XPA (Figure 1D-G), we further compared XPG and TFIIH association with DNA damage by FRAP after depletion of XPA. In cells depleted of XPA, XPD-mClover showed a slightly smaller UV-dependent immobile fraction than in siCTRL-treated cells (Figure 2C). This is similar to what we have previously observed in FRAP experiments with TFIIH subunit XPB in the absence of XPA, and suggests that XPA merely stabilizes the binding of TFIIH to DNA damage (59), rather than being absolutely required for its recruitment and binding. In contrast, the UV-induced XPG-mClover immobile fraction was much more diminished in XPA-depleted cells (Figure 2D), indicating that the association of TFIIH and XPG with DNA damage differently depends on XPA.

**Figure 2.**
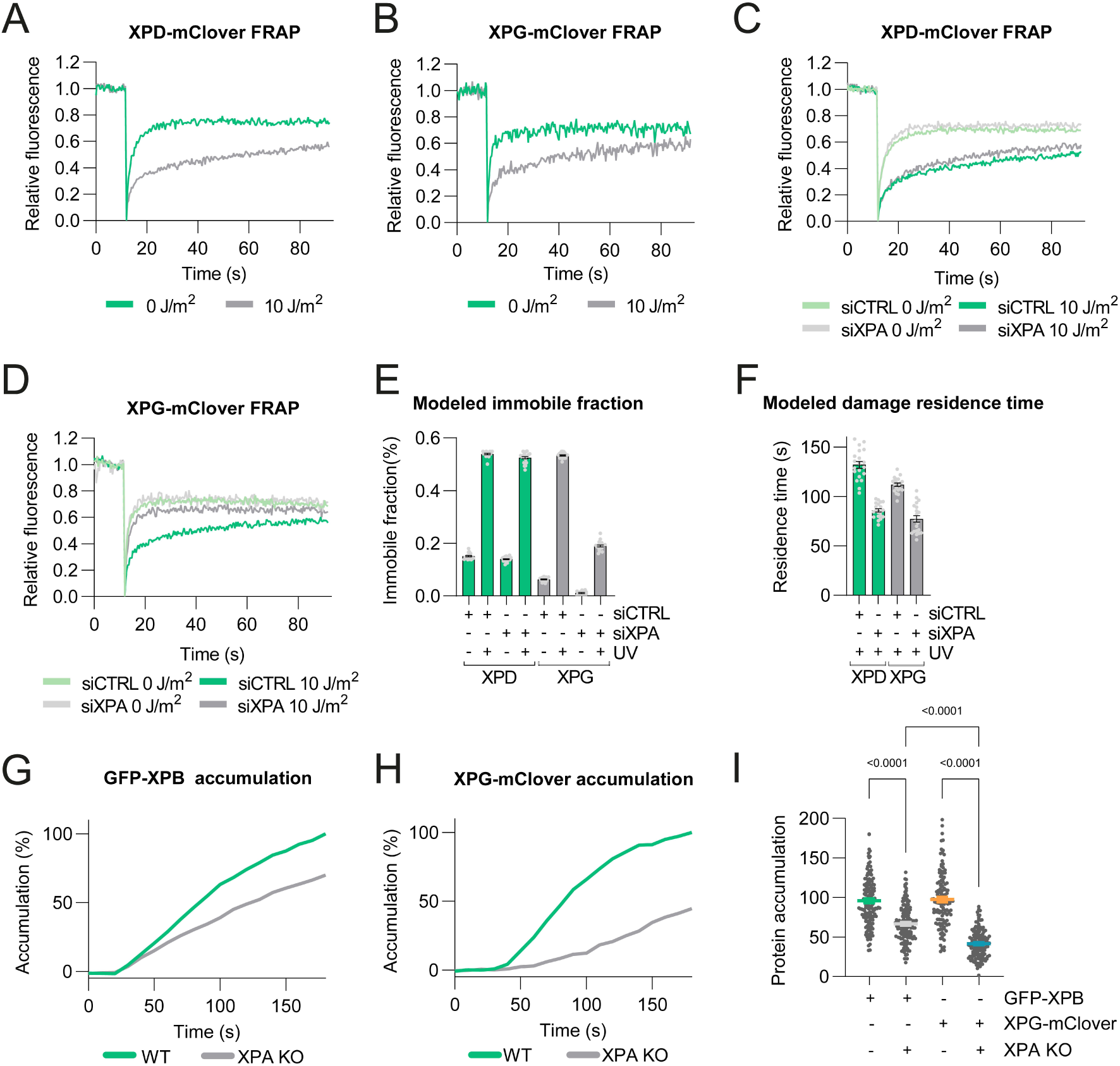
TFIIH and XPG association with DNA damage differently depends on XPA. (**A**) FRAP analysis of endogenous XPD mobility in U2OS XPD-mClover cells before (0 J/m^2^) and immediately after UV irradiation (10 J/m^2^). Number of cells: n= 30, 31 (0 and 10 J/m^2^). (**B**) FRAP analysis of endogenous XPG mobility in U2OS DIVa XPG-mClover cells before (0 J/m^2^) and immediately after UV irradiation (10 J/m^2^). Number of cells: n= 29, 30 (0 and 10 J/m^2^). (**C**) FRAP analysis of endogenous XPD mobility in U2OS XPD-mClover cells treated with control (CTRL) or XPA siRNA, non-irradiated (0 J/m^2^) or immediately after UV irradiation (10 J/m^2^). Number of cells: n= 30, 30 (for siCTRL 0 and 10 J/m^2^); n= 30, 30 (for siXPA 0 and 10 J/m^2^). (**D**) FRAP analysis of endogenous XPG mobility in U2OS DIVa XPG-mClover cells. Data from Figure 1F shown here for comparison with XPD FRAP. For all curves, XPD-mClover and XPG-mClover fluorescence recovery was measured in a strip across the nucleus after bleaching and normalized to the bleach depth. Each FRAP curve represents the average of three independent experiments. (**E**) Immobile fractions of XPD-mClover and XPG-mClover obtained from Monte Carlo simulations of FRAP data shown in C and D. Mean and SEM of the 20 best fitting simulations. (**F**) Residence time of XPD-mClover and XPG-mClover at UV-induced DNA damage obtained from Monte Carlo simulations of FRAP data shown in C and D. Mean and SEM of the 20 best fitting simulations. (**G**) Real-time imaging of GFP-XPB and (**H**) XPG-mClover accumulation at 266 nm UV-C laser-induced LUD in XPA proficient (WT) and XPA KO U2OS cells. Protein accumulation was normalized to pre-damage fluorescence. Each curve represents the mean of two independent experiments. (I) Protein accumulation at LUD calculated from the UV-C accumulation (determined between 160 and 180 s after UV-C irradiation) analyses depicted in G and H. Numbers in the graph represent *p-values* determined by ONE-WAY ANOVA. Mean and SEM of two independent experiments. Number of cells: n= 162, 161 (GFP-XPB and GFP-XPB XPA KO); n= 134, 148 (XPG-mClover and XPG-mClover XPA KO).

To test if the binding of TFIIH and XPG to DNA is indeed different, we used computational modeling to calculate and compare the XPG and XPD assembly and disassembly kinetics based on these FRAP experiments. To this end, we fitted the experimental FRAP curves to simulated curves obtained by Monte Carlo simulation (57, 65), to obtain average immobile fractions and residence times at DNA damage. Monte Carlo simulation suggested that in non-irradiated conditions, a higher fraction of XPD molecules is bound to chromatin, than XPG molecules (Figure 2E). This higher immobilized fraction of XPD molecules in undamaged cells reflects the specific activity of TFIIH in transcription initiation, as was previously also found with FRAP on ectopically expressed XPB-GFP (66). Strikingly, after UV irradiation, XPD and XPG showed similar immobile fractions (Figure 2E), likely reflecting their simultaneous activity in NER, but dissimilar residence times at DNA damage (Figure 2F). Following XPA depletion in UV-irradiated cells, only the immobile fraction of XPG molecules was significantly reduced (Figure 2E), while the residence times of both XPD and XPG at DNA damage were reduced. These simulated data therefore confirm that XPA activity differently affects TFIIH and XPG association with DNA damage. Also, this modeling challenges the idea that XPG localizes to DNA damage as part of a preformed TFIIH-XPG complex To independently confirm that XPG depends more on XPA than does TFIIH, with regard to DNA damage binding, we monitored XPG and TFIIH real-time recruitment to UV-C laser (266 nm)-induced DNA damage in XPA proficient and XPA deficient cells by live cell imaging. As alternative to the U2OS DIvA cells so far used, we used CRISPR/Cas9 to knock in mClover at the C-terminus of XPG in normal U2OS cells, and subsequently knocked out (KO) XPA in these cells (Supplementary Figure 1E). Also, we used previously generated GFP-XPB KI U2OS cells (59), as alternative to the XPD-mClover KI cells, and knocked out XPA in these cells as well (Supplementary Figure 1F). Both GFP-XPB and XPG-mClover were rapidly recruited to DNA damage in XPA proficient cells, which was diminished in XPA KO cells (Figure 2G-I). However, XPA loss had a different and stronger effect on XPG-mClover compared to GFP-XPB, causing mildly diminished GFP-XPB recruitment but severely delayed XPG-mClover recruitment to DNA damage (Figure 2H). We obtained even more pronounced results in cells treated with siRNA against XPA (Supplementary Figure 1G-I). These results are consistent with the milder XPA effects on TFIIH binding to DNA damage observed in FRAP (Figure 2C-F).

To discard the possibility that the residual XPG-mClover recruitment observed in XPA KO and siXPA-treated cells is due to the induction of lesions that are not repaired by NER, or due to the generation of R loops, to which XPG may localize, we knocked out XPC in XPG-mClover KI U2OS cells (Supplementary Figure 1E). In the absence of XPC, no accumulation of XPG-mClover at UV-C laser-induced DNA damage was detected (Supplementary Figure 1J, K), demonstrating that XPG-mClover accumulation solely reflects its activity in GG-NER. As comparison, we also generated XPA proficient and XPA KO U2OS cells with TEV-mAID-mClover knocked-in at the C-terminus of XPF (referred to as XPF-mClover) (Supplementary Figure 1L, M). Like XPG, XPF-mClover was rapidly recruited to laser-induced DNA damage, but this was, in contrast to XPG and TFIIH, completely XPA dependent (Supplementary Figure 1N, O), in line with our previous observations and those of others (36, 42, 43, 67). Thus, our UV-C laser accumulation experiments reveal that XPG depends more on XPA than does TFIIH to be recruited to NER lesions. Together, our data therefore indicate that TFIIH and XPG diffuse and localize to UV-induced DNA damage separately. In the absence of XPA, however, XPG is still able to localize to DNA damage, albeit less efficient, in contrast to ERCC1-XPF that is completely dependent on XPA.

### XPG residence time at DNA damage depends on XPF and PCNA

Once XPG and ERCC1-XPF are bound to damaged DNA, ERCC1-XPF is thought to first incise DNA 5’ to the lesion, after which XPG incises 3’ of the lesion. Both *in vitro* and *in vivo* partial DNA synthesis have been observed in the presence of catalytically active ERCC1-XPF but catalytically inactive XPG. This indicates that the 5’ incision is sufficient to initiate DNA repair synthesis, probably by already allowing PCNA and polymerase loading, whereas the 3’ incision is required to complete but not to initiate repair synthesis (44). It is still unclear whether XPG dissociates directly upon incision, or whether it might sometimes prematurely dissociate even if there is no incision, and whether it has a role in recruiting downstream factors. One such downstream factor could be PCNA, whose *in vitro* interaction with XPG, via a PIP box motif within XPG, has been reported (14). To better understand XPG dissociation, we measured and compared XPG-mClover, XPD-mClover and XPF-mClover residence times at DNA damage by inverse fluorescence recovery after photobleaching (iFRAP) in U2OS cells. To this end, we irradiated cells locally through a microporous filter to induce fluorescent protein accumulation at LUD, after which we monitored the fluorescence decrease in the LUD, while continuously bleaching the fluorescently tagged proteins outside the LUD (23).

iFRAP showed that XPG-mClover dissociation is slower than that of XPF-mClover but faster than that of XPD-mClover (Figure 3A and B). Possibly, the TFIIH residence time at DNA lesions is longer than that of XPF and XPG because TFIIH arrives earlier and interacts with multiple NER proteins, including XPC, XPA, XPG and XPF for the assembly of the initiation complex. The faster XPF dissociation, on the other hand, is confirmed by FRAP, which showed that after UV irradiation only a small fraction of XPF-mClover molecules transiently bound to DNA damage (Supplementary Figure 2A and B). This faster dissociation of XPF-mClover is in line with the idea that ERCC1-XPF cuts before XPG and therefore needs to bind DNA for a shorter time. The longer residence time of XPG at damaged DNA might indicate that XPG arrives before ERCC1-XPF and/or remains longer bound. Possibly, this indicates that XPG, before or after 3’ incision, helps in recruiting other factors, as also previous *in vitro* data have suggested (50). Recently, the translocase HLTF was shown to promote the dissociation of TFIIH and XPG after the dual incision by ERCC1-XPF and XPG (48). To study if the arrival of additional downstream factors affects XPG dissociation *in vivo,* we performed iFRAP after knocking down XPF (siXPF), RFC1 (siRFC1) and PCNA (siPCNA) by siRNA in XPG-mClover cells (Supplementary Figure 2C, D and E). XPG-mClover clearly dissociated slower upon XPF and PCNA depletion, but was unaffected by RFC1 depletion (Figure 3C and D). These results suggest that XPG dissociation depends on XPF-mediated incision, which allows subsequent incision by XPG, and therefore likely promotes its dissociation. Also, these results suggest that XPG dissociation is stimulated by the binding of PCNA, indicating that XPG interacts with PCNA and aids its recruitment after which it dissociates.

**Figure 3.**
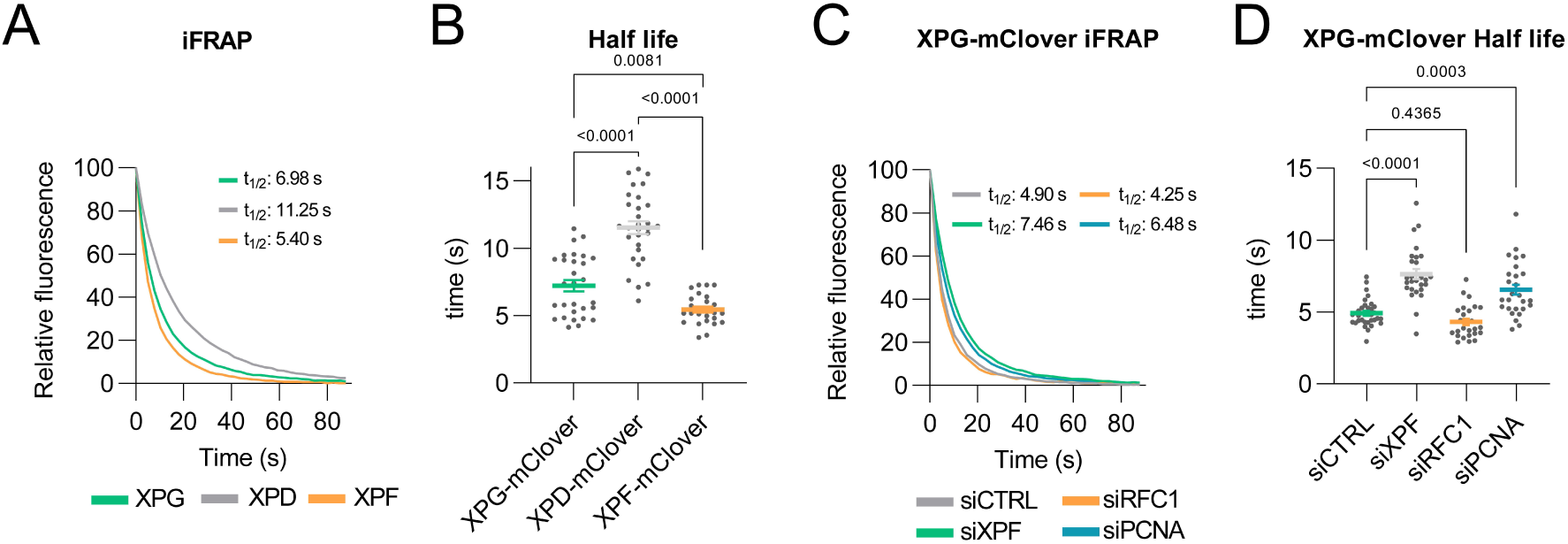
XPG residence at DNA damage is dependent on XPF 5’ incision and PCNA arrival. (**A**) iFRAP analyses reflecting XPG-mClover, XPD-mClover, and XPF-mClover dissociation from LUD in U2OS cells. Loss of fluorescence was measured over time and normalized to the intensity before bleaching. Each curve represents the mean of three independent experiments. Half-life for each condition is indicated. (**B**) Half-life of XPG-mClover, XPD-mClover, and XPF-mClover in the LUD as calculated from the iFRAP analyses depicted in A. Numbers in the graph represent *p-values* determined by ONE-WAY ANOVA. Mean and SEM of three independent experiments. Number of cells: n= 29 for XPG-mClover, n= 30 for XPD-mClover and n=26 for XPF-mClover. (**C**) iFRAP analyses reflecting XPG-mClover dissociation from LUD in U2OS cells, treated with control (CTRL), XPF, RFC1 or PCNA siRNA. Loss of fluorescence was measured over time and normalized to the intensity before bleaching. Each curve represents the mean of three independent experiments. Half-life for each condition is indicated (**D**) Half-life of XPG-mClover in the LUD as calculated from the iFRAP analyses depicted in C. Numbers in the graph represent *p-values* determined by ONE-WAY ANOVA. Mean and SEM of three independent experiments. Number of cells: n=31 for siCTRL, n=27 for siXPF, n=26 for siRFC1, n=28 for siPCNA.

### XPG incision of DNA promotes its dissociation

To test if XPG’s own catalytic activity is a prerequisite for its dissociation from DNA damage, we generated an XPG catalytically inactive mutant by introducing the E791A point mutation (16) in *XPG* in XPG-mClover KI U2OS cells using CRISPR/Cas9 technology (Figure 4A). We tested the UV sensitivity of mutant XPG(E791A)-mClover KI cells in a UV-colony survival assay, which confirmed that these cells are NER deficient due to the lack of XPG’s catalytic activity (Figure 4B). Next, we performed FRAP experiments to assess if XPG(E791A) dynamics and binding to damaged chromatin differ from those of wild type (WT) XPG. In non-treated cells, both XPG-mClover and XPG(E791A)-mClover showed similar mobility in FRAP, indicating that WT and mutant XPG molecules both freely diffuse through the nucleus. Upon UV irradiation, however, XPG(E791A)-mClover showed a much stronger immobilization than WT XPG-mClover (Figure 4C). Fitting these FRAP curves to Monte Carlo simulated curves (57, 65) indicated that not the fraction of bound XPG molecules, but the residence time of E791A mutant XPG at DNA damage is increased compared to WT XPG (Figure 4D, E). To test this experimentally, we used iFRAP, which showed that indeed the residence time of XPG(E791A)-mClover at damaged DNA is longer than that of WT XPG-mClover (Figure 4F, G). Thus, these results demonstrate that XPG dissociation is strongly promoted by its catalytic activity.

**Figure 4.**
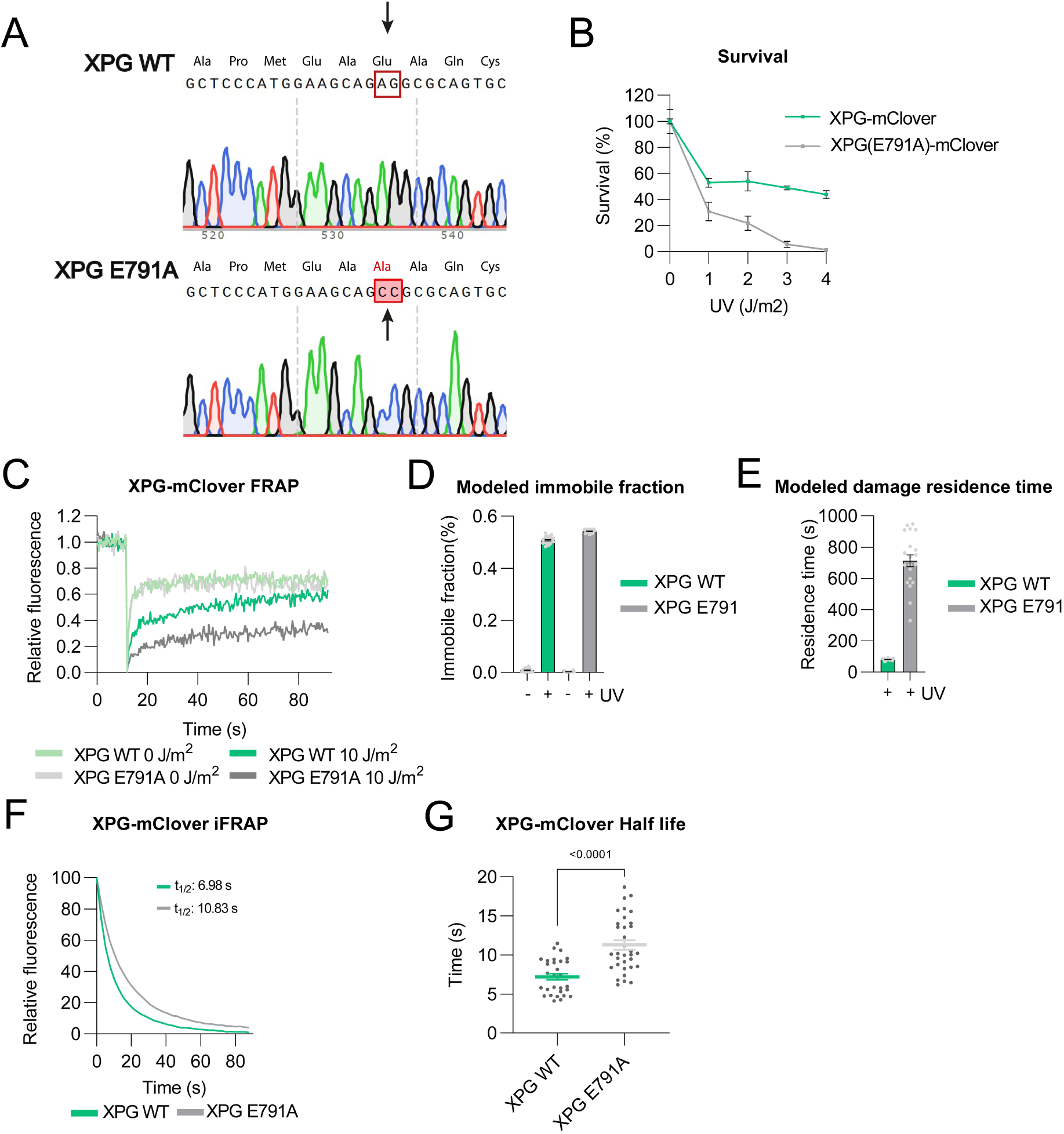
Catalytically inactive XPG resides longer at DNA damage. (**A**) DNA sequences of XPG encoding amino acids 786 to 794. Upper sequence depicts XPG WT DNA sequence with amino acids. Lower sequence depicts the E791A mutation, which changes the amino acid from Glu > Ala, as found by sequencing in our XPG(E791)-mClover mutant knock-in cells. (**B**) UV-C colony survival of U2OS DIVa XPG-mClover knock-in and U2OS DIVa XPG(E791A)-mClover knock-in cells after UV irradiation at the indicated doses. Survival was plotted as the percentage of colonies obtained after treatment normalized to the mean number of colonies in untreated conditions, set as 100%. Mean & SEM of three independent experiments, each performed in triplicate. (**C**) FRAP analysis of wild type (WT) and mutant XPG mobility in XPG-mClover and XPG(E791A)-mClover U2OS DIVa cells, before (0 J/m^2^) and immediately after UV irradiation (10 J/m^2^). FRAP curve of XPG-mClover from Figure 2B shown here for comparison with XPG(E791A) FRAP. Each curve represents the mean of three independent experiments. Number of U2OS DIVa XPG(E791A)-mClover cells: n= 28, 30 (0 and 10 J/m^2^). (**D**) Immobile fractions of XPG-mClover and XPG(E791A)-mClover obtained from Monte Carlo simulations of FRAP data shown in C. Mean and SEM of the 20 best fitting simulations. (**E**) Residence time of XPG-mClover and XPG(E791A)-mClover at UV-induced DNA damage obtained from Monte Carlo simulations of FRAP data shown in C. Mean and SEM of the 20 best fitting simulations. (**F**) iFRAP analyses reflecting XPG-mClover and XPG(E791A)-mClover dissociation from LUD in U2OS DIVa cells. Loss of fluorescence was measured over time and normalized to the intensity before bleaching. iFRAP curve of XPG-mClover from Figure 3A shown here for comparison with XPG(E791A) iFRAP. Each curve represents the mean of three independent experiments. Half-life for each condition is indicated (**G**) Half-life of XPG-mClover and XPG(E791A)-mClover in the LUD as calculated from the iFRAP analyses depicted in E. Numbers in the graph represent *p-values* determined by ONE-WAY ANOVA. Mean and SEM of three independent experiments. Number of U2OS DIVa XPG(E791A)-mClover cells: n= 33.

### EXO1 promotes XPG dissociation

Even though XPG(E791A)-mClover resides longer at DNA damage, it is still able to dissociate. This dissociation may be spontaneous or could involve other proteins that actively promote XPG displacement. We, therefore, tested involvement of the 5’-3’ exonuclease EXO1, which has been suggested to compete with canonical DNA repair synthesis factors for gap processing at problematic lesions and to generate long ssDNA intermediates that induce checkpoint activation (52, 53). In case of catalytically inactive XPG, the absence of 3’ incision and slow dissociation of XPG impedes efficient refilling during repair, which could allow EXO1 loading and DNA resection in a 5’ to 3’ direction leading also to the removal of stalled XPG from the lesion. Using immunofluorescence to visualize endogenously expressed EXO1, we confirmed that EXO1 is indeed recruited to UV lesions (Supplementary Figure 2F).

We, therefore, studied if the depletion of EXO1 affected XPG-mClover localization to DNA damage by immunofluorescence, UV-C laser accumulation and iFRAP experiments. As control, we depleted XPF. Depletion of EXO1 led to increased recruitment of WT XPG-mClover to LUD, similar as after XPF depletion, which was especially clear at 40 min after UV irradiation in immunofluorescence (Figure 5A, Supplementary Figure 2G). Recruitment of XPG(E791A)-mClover was higher than that of WT XPG, but also this was increased after depletion of XPF or EXO1. After DNA damage infliction by UV-C laser, recruitment of both WT XPG-mClover and XPG(E791A)-mClover was increased to the same level after depletion of XPF (Figure 5B and C). Strikingly, after EXO1 depletion XPG accumulation also increased, but this was even stronger for XPG(E791A)-mClover, suggesting that if NER cannot proceed properly, EXO1’s function may become even more important in processing the damaged DNA and promoting the dissociation of XPG. iFRAP confirmed this delayed dissociation of XPG-mClover from LUD in EXO1-depleted cells, which was even more pronounced for XPG(E791A)-mClover (Figure 5D and E).

**Figure 5.**
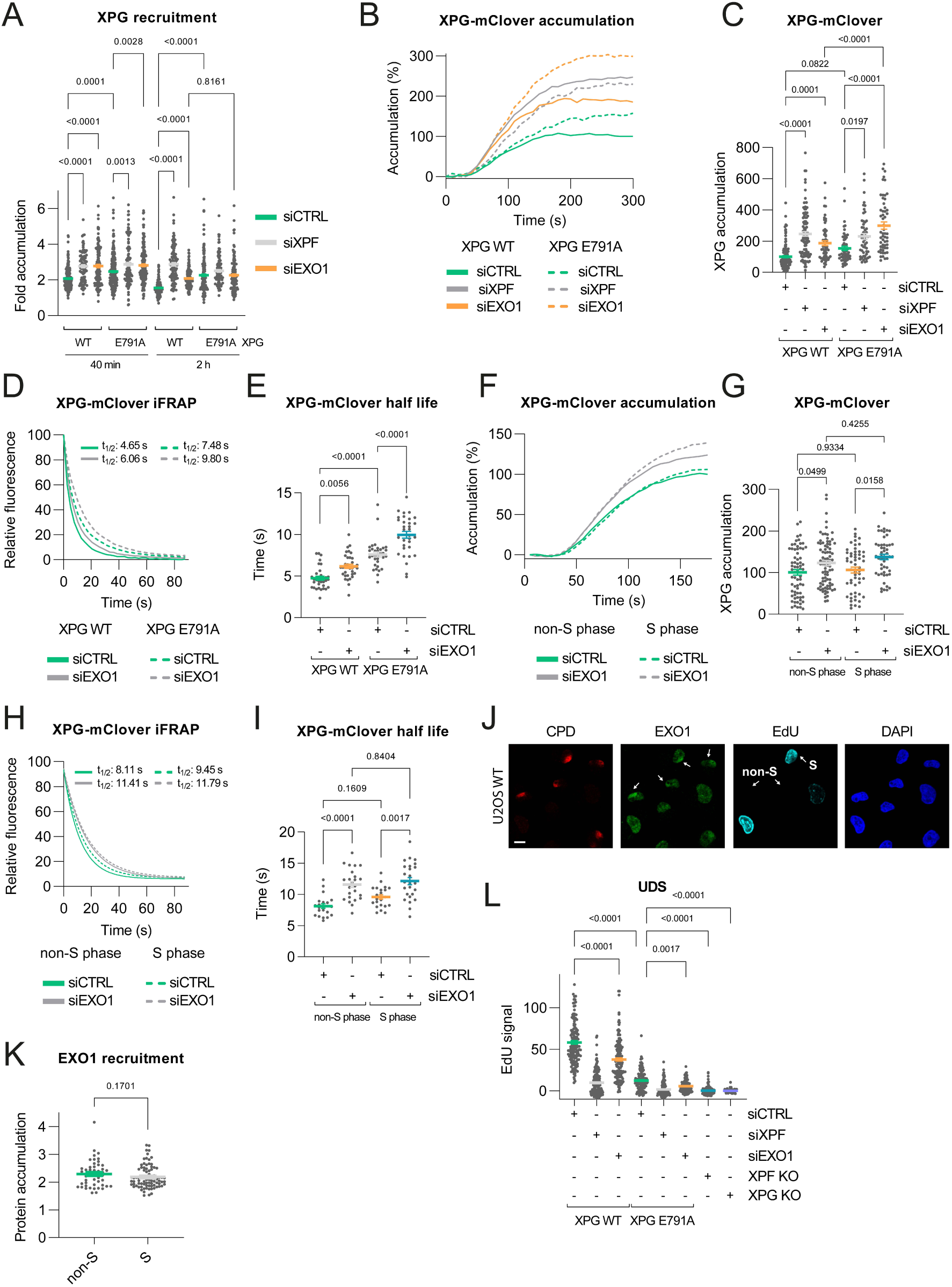
EXO1 promotes dissociation of XPG from DNA damage. (**A**) Quantification of XPG recruitment at LUD in immunofluorescence of XPG-mClover and XPG(E791A)-mClover U2OS DIVa cells, treated with control (CTRL), XPF or EXO1 siRNA, 40 min or 2 h after UV irradiation with 60 J/m^2^ UV-C through a microporous filter. The fold accumulation was calculated by normalizing fluorescence intensity at sites of LUD to the nuclear background and plotted as the average of the number of cells per condition from three independent experiments. Number of cells: n= 219, 93, 134 (for 40 min XPG WT siCTRL, siXPF, siEXO1); n=193, 111, 156 (for 2 h XPG WT siCTRL, siXPF, siEXO1); n= 237, 122, 173 (for 40 min XPG E791A siCTRL, siXPF, siEXO1); n= 197, 113, 143 (for 2 h XPG WT siCTRL, siXPF, siEXO1). Numbers in the quantification graphs represent *p-values* determined by ONE-WAY ANOVA. Error bars represent SEM. (**B**) Real-time imaging of XPG-mClover (WT) and XPG(E791A)-mClover accumulation at 266 nm UV-C laser-induced LUD in cells treated with control (CTRL), XPF or EXO1 siRNA. Protein accumulation was normalized to pre-damage fluorescence. Each curve represents the mean of two independent experiments. (**C**) XPG accumulation in the LUD calculated from the UV-C accumulation (determined between 280 and 300 s after UV-C irradiation) analyses depicted in (B). Numbers in the graph represent *p-values* determined by ONE-WAY ANOVA. Mean and SEM of two independent experiments. Number of cells: n= 136, 101, 59 (for XPG WT siCTRL, siXPF, and siEXO1); n= 52, 54, 56 (for XPG(E791A) siCTRL, siXPF, and siEXO1). (**D**) iFRAP analyses reflecting XPG-mClover (WT) and XPG(E791A)-mClover dissociation from LUD in U2OS DIVa cells treated with CTRL and EXO1 siRNA. Loss of fluorescence was measured over time and normalized to the intensity before bleaching. Each curve represents the mean of three independent experiments. Half-life for each condition is indicated (**E**) Half-life of XPG-mClover (WT) and XPG(E791A)-mClover in the LUD as calculated from the iFRAP analyses depicted in (D). Numbers in the graph represent *p-values* determined by ONE-WAY ANOVA. Mean and SEM of three independent experiments. Number of cells: n= 33, 35 (for XPG WT siCTRL, siEXO1); n= 33, 32 (for XPG E791A siCTRL, siEXO1). (**F**) Real-time imaging of XPG-mClover accumulation at 266 nm UV-C laser induced LUD in non-S and S phase cells in U2OS DIVa XPG-mClover mScarlet-PCNA knock-in cells treated with CTRL or EXO1 siRNA. XPG accumulation was normalized to pre-damage fluorescence. Each curve represents the mean of three independent experiments. (**G**) XPG accumulation in the LUD calculated from the UV-C accumulation (determined between 228 and 243 s after UV-C irradiation) analyses depicted in (F). Numbers in the graph represent *p-values* determined by ONE-WAY ANOVA. Mean and SEM of three independent experiments. Number of cells: n= 68, 85 (for non-S phase siCTRL and siEXO1); n= 55, 51 (for S-phase siCTRL and siEXO1). (**H**) iFRAP analyses reflecting XPG-mClover dissociation from LUD in non-S and S phase cells in U2OS DIVa cells treated with CTRL or EXO1 siRNA. Loss of fluorescence was measured over time and normalized to the intensity before bleaching. Each curve represents the mean of three independent experiments. Half-life for each condition is indicated (**I**) Half-life of XPG-mClover in the LUD as calculated from the iFRAP analyses depicted in (H). Numbers in the graph represent *p-values* determined by ONE-WAY ANOVA. Mean and SEM of three independent experiments. Number of cells: n= 21, 24 (for non-S phase siCTRL and siEXO1); n= 22, 25 (for S-phase siCTRL and siEXO1). (**J**) Representative immunofluorescence images of U2OS WT cells 30 min after UV irradiation with 60 J/m^2^ UV-C through a microporous filter to induce LUD. Images show accumulation of endogenous EXO1 at LUD in non-S and S phase cells as distinguished by EdU staining. Coverslips were stained with antibodies against EXO1 and CPD. DNA is stained with DAPI. Scale bar 10 µm. (**K**) Quantification of EXO1 recruitment to LUD in non-S and S-phase cells as described in (J). The fold accumulation was calculated by normalizing fluorescence intensity at sites of local damage to the nuclear background and plotted as the average of the number of cells per condition from two independent experiments. Number of cells: n= 49 for non-S: n= 83 for S cells. (**L**) Quantification of Unscheduled DNA synthesis (UDS) in U2OS DIVa XPG-mClover (WT), U2OS DIVa XPG(E791A)-mClover, XPF KO and XPG KO U2OS cells, untreated or treated with CTRL, XPF, or EXO1 siRNA. UDS was determined after EdU incorporation for 3 h following 60 J/m^2^ irradiation through a microporous filter. EdU fluorescence levels were quantified after background subtraction. Numbers in the graph represent *p-values* determined by ONE-WAY ANOVA. Mean & SEM of three independent experiments. Number of cells: n= 152, 152, 145 (for XPG WT siCTRL, siXPF and siEXO1); n= 152, 152, 151 (for XPG E791A siCTRL, siXPF and siEXO1); n= 188 for XPF KO; n= 151 for XPG KO.

Since the cell cycle phase is thought to be an important determinant of which DNA polymerases are used for DNA synthesis during NER (51) and in previous studies EXO1 recruitment to UV-induced DNA lesions was only observed in non-S phase cells (52, 53), we tested whether EXO1-stimulated XPG dissociation is different depending on the cell cycle. Hence, we inserted an mScarlet tag at the N-terminus of PCNA in XPG-mClover KI U2OS cells, which allowed us to differentiate between non-S and S phase cells by discriminating replication-associated PCNA foci (68) (Supplementary Figure 2H and I). UV-C laser accumulation experiments showed that XPG-mClover accumulation is similar in both S and non-S phase cells and increases after EXO1 depletion irrespective of the cell cycle phase (Figure 5F and G). iFRAP confirmed that XPG-mClover dissociation from sites of DNA damage was delayed in EXO1-depleted cells, independently of the cell cycle phase (Figure 5H and I). Together, these data show that XPG dissociation from DNA damage is stimulated by EXO1 and that EXO1 especially promotes the removal of catalytically inactive XPG when 3’ incision does not occur.

As EXO1 depletion affects XPG dissociation in both S- and non-S phase cells, we tested whether EXO1 is similarly recruited to LUD in non-S and S cells. To this end, we incubated U2OS cells with 5-ethynyl-2-deoxiuridine (EdU) to distinguish replicating cells by fluorescently visualizing EdU incorporated into DNA. Subsequent immunofluorescence studies showed that EXO1 is clearly recruited to UV-induced lesions, as indicated by CPD staining, in both S phase and non-S phase cells (Figure 5J and K), in line with the fact that we did not observe a cell cycle-dependent effect on XPG release upon EXO1 depletion (Figure 5H and I). We finally tested if the lack of EXO1 also results in reduced UV-induced DNA repair synthesis, referred to as unscheduled DNA synthesis (UDS), as EXO1 is thought to compete with canonical gap filling during NER by extending the DNA gap, which would lead to more DNA synthesis (54). We, therefore, tested if EXO1 depletion affected UDS in XPG WT and XPG(E791A) cells by measuring EdU incorporation in LUD. As control, we used previously generated XPF and XPG KO cells (59), which showed that, as expected, the UV-dependent UDS signal is completely dependent on the presence of both NER endonucleases (Figure 5L). In WT cells, a significant part of the UV-induced DNA repair synthesis also depended on EXO1. Moreover, we observed that the UDS signal was severely reduced in XPG(E791A) cells, showing that most UV-dependent DNA synthesis is dependent on XPG catalytic activity. However, a residual UDS signal was still detected in XPG(E791A) cells, as previously also reported (44), which was reduced upon depletion of XPF and EXO1. These data therefore suggest that EXO1 can compete with canonical NER gap processing in cells both proficient and deficient in XPG catalytic activity. As XPF incises DNA when catalytically active or inactive XPG is present, but not when XPG is completely absent (44), our results suggest that EXO1 can bind to DNA after the 5’ incision by XPF, and can compete with XPG to start extensive DNA resection while XPG is displaced.

## Discussion

In this study, we show that the stable assembly of XPG into the NER incision complex is stimulated by both TFIIH and XPA, but that only TFIIH is essential for XPG recruitment. Our data suggest that, although XPG can form a complex with TFIIH in the absence of DNA damage (46), XPG localizes separately from TFIIH to DNA damage. Moreover, we show that XPG resides longer than XPF at DNA damage and that its dissociation is stimulated by its own incision activity, by that of XPF, and by the subsequent recruitment of PCNA. These results suggest that XPG dissociates later than XPF to help to recruit downstream factors. Moreover, we show that EXO1 is recruited to UV lesions and promotes XPG dissociation, even when XPG itself does not incise DNA. We therefore propose, in line with previous research (52, 54, 69), that EXO1 can resect the XPF-incision-generated 5’ flap DNA structure in a 5’-3’ direction, which leads to XPG removal from the damaged DNA.

In agreement with previous immunofluorescence experiments in XPD and XPB deficient fibroblasts (47), we show that XPG recruitment requires TFIIH. Co-immunoprecipitation experiments have shown that XPG interacts with and stabilizes TFIIH, even in the absence of DNA damage, forming a stable TFIIH-XPG complex that is active in *in vitro* transcription and DNA repair assays (46). We therefore addressed whether in intact cells, XPG is already bound to TFIIH upon engagement in NER. We provide several lines of evidence to show that XPG associates with TFIIH at the site of damage. First, FRAP data of endogenously-tagged XPG and XPD proteins, analyzed by Monte Carlo simulations, showed different immobile fractions in unperturbed cells, which only became more similar after UV irradiation. Also, we observed a different dependency on XPA for TFIIH and XPG binding to damaged DNA. In line with these results, previous FRAP studies on XPG-eGFP and XPB-eGFP, overexpressed in Chinese hamster ovary cells and human patient fibroblasts, already showed that, in unperturbed conditions, XPG freely diffuses in the nucleus and that its mobility was temperature independent, while that of TFIIH varied at different temperatures (47, 66). Moreover, we find that the dissociation of XPG at sites of damage is faster than that of TFIIH, indicating that XPG is not part of the same associating or dissociating TFIIH complex.

Even if TFIIH and XPG assemble sequentially in the NER incision complex, it is likely that TFIIH and XPG interact before the DNA bubble generated by TFIIH is fully completed. *In vitro* studies revealed that XPG strongly stimulates XPD helicase activity, possibly by stabilizing its binding to DNA and promoting its transition from ssDNA translocation to dsDNA unwinding at the ss/dsDNA border (33, 70). Structural modeling of the NER incision complex suggested that XPG inserts a hydrophobic wedge and β-hairpin motif at the 3’ ss/dsDNA border that could act as ‘helicase pin’ to facilitate efficient strand separation by XPD activity (71). Interestingly, XPG was shown to incise DNA only when TFIIH stalls at a lesion and a stretch of ssDNA of a certain length is formed (70), implying that XPG stimulates TFIIH DNA unwinding activity and that in turn TFIIH stimulates XPG incision activity. Upon its recruitment to DNA damage, XPG is thought to exchange with XPC, as inefficient XPC dissociation impairs stable association of XPG with the incision complex (72, 73). In fact, XPC and XPG, as well as their yeast orthologs Rad4 and Rad2, respectively, share similar acidic binding motifs with which they interact with the PH domain of p62 (21, 71, 74), leading to mutual exclusion when bound to TFIIH. Therefore, it may be that, upon DNA damage detection, TFIIH first forms a stable complex together with XPC, which anchors TFIIH to DNA damage and helps it to initiate DNA unwinding and DNA damage verification (23, 30, 75). Subsequently, before DNA unwinding by TFIIH is finished, XPC is replaced by XPG. Then, XPG, together with XPA, stimulates the XPD helicase to complete the generation of the NER bubble and of the ss/dsDNA junction substrate at which XPG is positioned for 3’ incision. It will be of interest to investigate if only the physical presence of XPG or also the XPG catalytic activity is needed for stimulation of XPD helicase activity. Also, XPG has been reported to form homodimers (16, 76), but the exact functional and/or structural relevance of these in the build-up of the incision complex and XPG function in NER remains unknown.

Dual incision and DNA synthesis during NER have been shown to be tightly coordinated. XPF 5’ incision is triggered by the presence of XPG and this is followed by XPG 3’ incision and DNA synthesis. Interestingly, initiation of DNA synthesis was already detected after 5’ incision even in the absence of 3’ incision, whereas completed DNA synthesis was only observed after dual incision (44). We also observed low levels of XPF-dependent DNA synthesis (UDS) in our XPG-E791A mutant KI cells. However, these levels appeared to be lower than previously observed in XPG deficient XPCS1RO fibroblasts overexpressing E791A-XPG mutant cDNA, which may be because of the lower endogenous XPG levels in our cells. This residual DNA synthesis suggests that XPF dissociates after its 5’ incision, and can even do so in absence of XPG catalytic activity, to allow loading of PCNA and DNA synthesis factors at the generated free 3’-OH end, 5’ to the lesion, to initiate repair synthesis, as was previously suggested (77). In support of this rapid XPF release, we show by FRAP and iFRAP that XPF resides for a shorter time at the damaged DNA than XPG and XPD/TFIIH. Moreover, the residence time of XPG increased in the absence of XPF, indicating that XPG dissociation is stimulated by XPF-mediated incision. The longer residence time of XPD might be because of the earlier arrival of TFIIH, although we cannot discard the possibility that its role in transcription influences these results. Furthermore, the longer residence time of XPG might indicate that this endonuclease does not incise and dissociate immediately upon XPF incision. Interestingly, we also show that XPG resides longer at the damage in absence of PCNA (Figure 3C, D), indicating that it may remain bound to DNA damage to help recruit PCNA through its PIP box, even before PCNA is stably loaded onto DNA by RFC, as was also shown in *in vitro* immobilized template assays (14, 50, 63, 78). We did not observe that XPG dissociation was affected by RFC depletion, which indicates that XPG has no role in its recruitment. RFC could be recruited by RPA, as suggested before (50), and is required to load PCNA. Our results also contemplate the possibility that XPG dissociates together with TFIIH (49) after eviction of DNA damage by the translocase HLTF (48). Besides direct protein-protein interactions, possibly also post-translational modifications may influence XPG assembly to and disassembly from the NER incision complex. For example, quantitative proteomic studies reported ubiquitylation of XPG after UV irradiation (79, 80). Although the role of this modification is not yet understood, it has been proposed that after 3’ incision the E3 ubiquitin ligase complex CRL4^Cdt2^ ubiquitylates XPG, targeting the protein for removal from the NER complex and degradation (81). It will be interesting to further investigate the role of such PTMs in regulating XPG activity in NER in the future.

In addition to XPF and PCNA, we show here that XPG dissociation is promoted by the exonuclease EXO1, whose implication in NER was thus far limited to few studies in yeast and mammalian cells. In these studies, it was shown that ssDNA gaps produced by NER activity after UV irradiation could be enlarged by DNA resection by the 5’-3’ exonuclease activity of EXO1 (53, 54). In line with this, we found a lower UDS signal upon EXO1 depletion, indicating that UV-induced DNA synthesis is partly dependent on EXO1 activity. It was proposed that this EXO1-processing of NER intermediates is restricted to a subset of UV-induced lesions, such as lesions for which DNA synthesis is somehow impeded. Such lesions could be closely spaced lesions on opposite DNA strands. Normally, for single lesions, DNA repair synthesis is thought to be faster than exonucleolytic processing by EXO1, preventing EXO1 to generate extended ssDNA intermediates (52). However, for lesions in close proximity on opposite DNA strands, the lesion in the template strand could impede gap filling by the regular DNA polymerases Polδ and Polɛ. EXO1 was proposed to be recruited after 5’-incision by XPF to process such NER intermediates (54, 69). Furthermore, alternatively to the regular DNA polymerases, the Y-family TLS polymerases Polκ, Polι, Polη, and REV1 were found to be recruited to gap intermediates processed by EXO1. These TLS polymerases, especially Polκ, were found to be able to synthesize DNA past lesions on the opposite strand, leading to removal of EXO1 from damaged DNA (52). This competition between canonical gap filling and exonucleolytic processing may also occur when XPG incision is delayed or cannot take place, i.e. when DNA repair synthesis is initiated but not readily completed, suggesting that EXO1 processing may also compete with XPG incision. Interestingly, by iFRAP we found that both enzymatically active and inactive XPG reside longer at sites of DNA damage in the absence of EXO1. This indeed suggests that EXO1 promotes the dissociation of XPG at certain lesions, likely by extending the ssDNA gap, which implies that XPG does not need to incise the DNA and is displaced. This was confirmed by our observation that the low residual UDS signal in XPG(E791A) mutant cells was dependent on both XPF and EXO1, indicating that in absence of XPG 3’ incision, EXO1 can still process the DNA. It will be interesting to verify this further by measuring XPG retention at sites of damage in cells in which EXO1 is still present but its catalytic activity impaired. Finally, we found that EXO1 function in NER acts independent of the cell cycle, which was supported by its recruitment to UV lesions in S and non-S phase cells.

In summary, we show that the recruitment and activity of XPG in NER is tightly regulated by both pre-incision and post-incision factors. To further clarify the exact role of EXO1 it will be necessary to study whether any NER-specific protein interactions are involved in EXO1 recruitment. Also, it will be interesting to determine which other proteins act together with EXO1 in dealing with lesions that are repaired in a manner that apparently does not require XPG’s catalytic activity.

## Acknowledgements

We thank Gert van Cappellen and Gert-Jan Kremers of the Erasmus MC Optical Imaging Center for microscope support. We also thank Marvin van Toorn for his helpful insights and Carla Engel for technical assistance. This work was supported by the Netherlands Organization for Scientific Research (ALWOP.494), Erasmus MC (HDMA Grant), the Dutch Cancer Society (KWF 10506), the European Research Council (advanced grant 340988-ERC-ID) Oncode Institute partly financed by Dutch Cancer Society.

## Author contribution

AMV, CRS and CDM performed live cell imaging and other human cell line experiments. AMV and MG performed UV survival experiments. KLT generated DNA constructs. AMV, AFT and JvG generated cell lines. BG and AH performed Monte Carlo simulations. JP supervised JvG. AMV, CRS, CDM, WV and HL conceptualized ideas, designed experiments and analyzed data. AMV and HL wrote the manuscript. All authors reviewed the manuscript.

