## Supplemental information for "Regulation of XPG-DNA damage binding dynamics by pre- and post-incision nucleotide excision repair factors and EXO1"

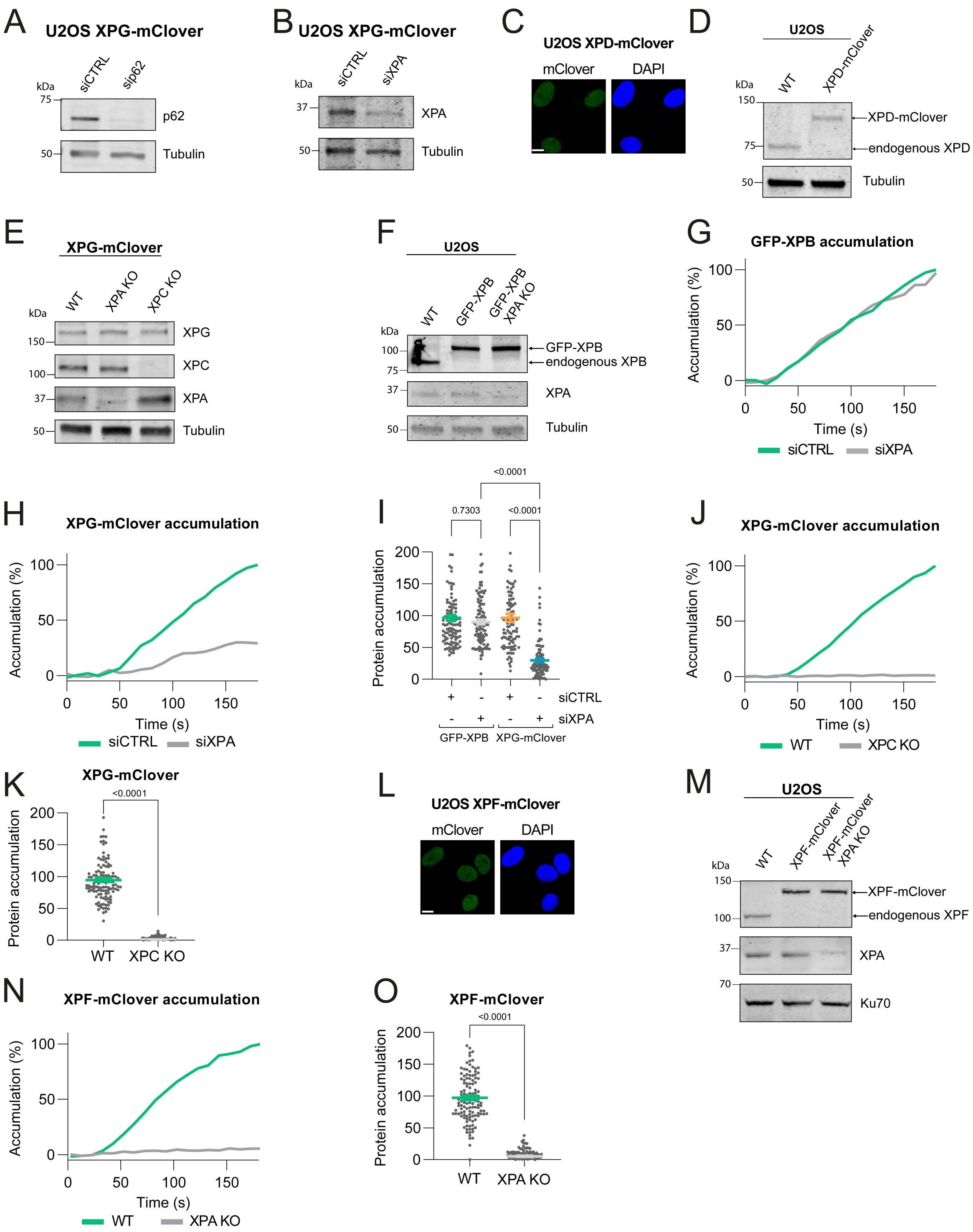


**Supplementary Figure S1.** Immunoblots, fluorescence images of knock-in cells and UV-C laser accumulation. (**A** and **B**) Immunoblot analyses of XPG-mClover U2OS DIVa cells treated with control, p62 or XPA siRNA. Immunoblots were stained with antibodies against p62, XPA and tubulin (loading control). (**C**) Representative image of fixed U2OS XPD-mClover knock-in cells. DNA was stained with DAPI. Scale bar represents 10 µM. (**D**) Immunoblot analysis of WT and XPD-mClover knock-in U2OS cells. Immunoblot was stained with antibodies against XPD and tubulin (loading control). (**E**) Immunoblot analysis of XPA knockout in U2OS XPG-mClover and XPC knockout in U2OS DIVa XPG-mClover cells. Immunoblot was stained with antibodies against XPG, XPC, XPA and tubulin (loading control). (**F**) Immunoblot analysis of WT and GFP-XPB knock-in U2OS cells either proficient or knockout for XPA. Immunoblot was stained with antibodies against XPB, XPA and tubulin (loading control). (**G** and **H**) Real-time imaging of GFP-XPB and XPG-mClover accumulation at 266 nm UV-C laser induced LUD in cells treated with control (CTRL) or XPA siRNA. Protein accumulation was normalized to pre-damage fluorescence. Each curve represents the mean of two independent experiments. (**I**) XPB and XPG accumulation in the LUD calculated from the UV-C accumulation irradiation (determined between 160 and 180 s after UV-C) analyses depicted in (G and H). Numbers in the graph represent *p-values* determined by

ONE-WAY ANOVA. Mean and SEM of two independent experiments. Number of cells: n= 110, 105 (for GFP-XPB siCTRL and siXPA); n= 99, 92 (for XPG-mClover siCTRL and siXPA). (**J**) Real-time imaging of XPG-mClover accumulation at 266 nm UV-C laser induced LUD in WT and XPC KO XPG-mClover knock-in U2OS DIVa cells. XPG accumulation was normalized to pre-damage fluorescence. Each curve represents the mean of three independent experiments. (**K**) XPG accumulation in the LUD calculated from the UV-C accumulation (determined between 160 and 180 s after UV-C irradiation) analyses depicted in (J). Numbers in the graph represent *p-values* determined by ONE-WAY ANOVA. Mean and SEM of three independent experiments. Number of cells: n= 120, 155 (for WT and XPC KO). (**L**) Representative image of fixed U2OS XPF-mClover knock-in cells. DNA was stained with DAPI. Scale bar represents 10 µM. (**M**) Immunoblot analysis of WT and XPF-mClover knock-in U2OS cells either proficient or knockout for XPA. Immunoblot was stained with antibodies against XPF, XPA and Ku70 (loading control). (**N**) Real-time imaging of XPF-mClover accumulation at 266 nm UV-C laser induced LUD in XPA proficient or XPA KO cells. Protein accumulation was normalized to pre-damage fluorescence. Each curve represents the mean of two independent experiments. (**O**) XPF accumulation in the LUD calculated from the UV-C accumulation (determined between 160 and 180 s after UV-C irradiation) analyses depicted in (N). Numbers in the graph represent *p-values* determined by ONE-WAY ANOVA. Mean and SEM of two independent experiments. Number of cells: n= 125, 108 (for WT and XPA KO).

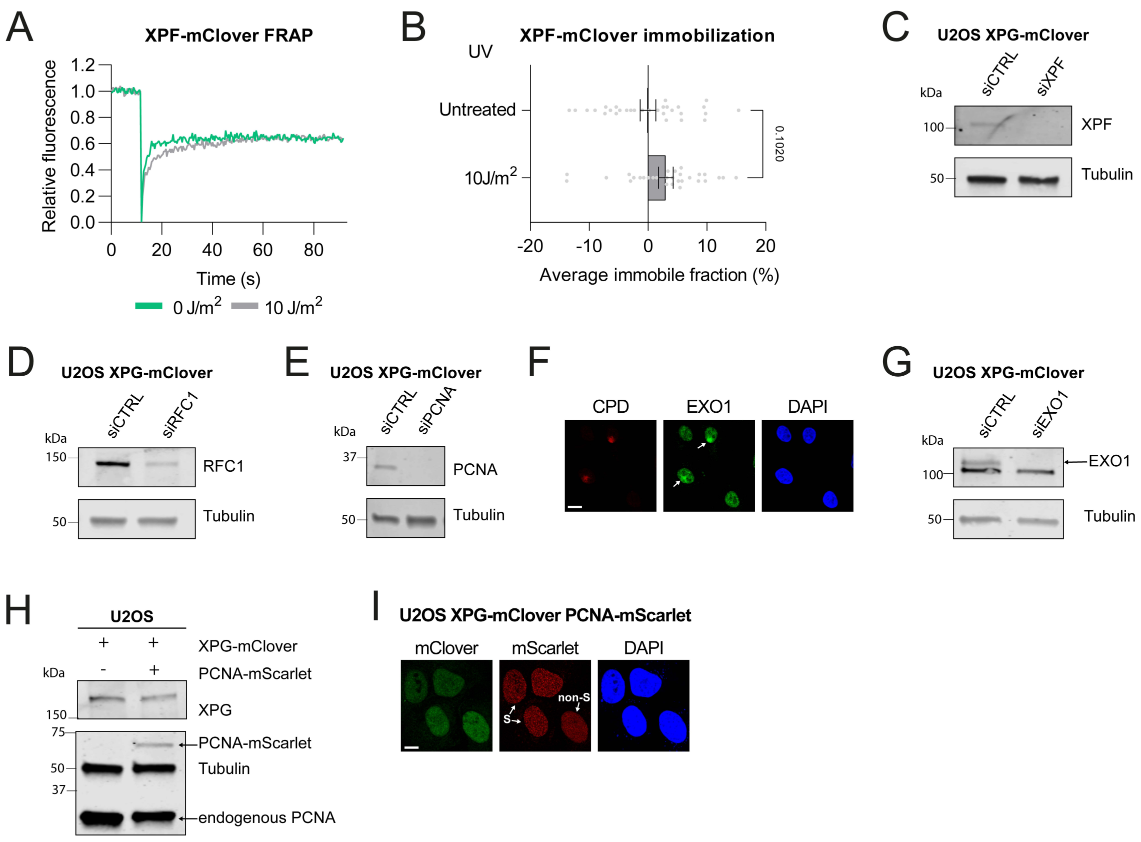


**Supplementary Figure S2.** FRAPs, immunoblots, immunofluorescence of knock-in cells and EXO1 recruitment to LUD. (**A** and **B**) FRAP analysis of endogenous XPF mobility in U2OS XPF-mClover cells and quantification of immobile fraction before (0 J/m$^2$) and immediately after UV irradiation (10 J/m$^2$). Number of cells: n= 30, 30 (for 0 and 10 J/m$^2$). For all curves, XPF-mClover fluorescence recovery was measured in a strip across the nucleus after bleaching and normalized to the bleach depth. Each FRAP curve represents the average of three independent experiments. Numbers in the quantification graph represent *p-values* determined by ONE-WAY ANOVA. Error bars represent SEM. (**C-E**) Immunoblot analyses of XPG-mClover U2OS DIVa cells treated with control, XPF, RFC1 or PCNA siRNA. Immunoblots were stained with antibodies against XPF, RFC1, PCNA and tubulin (loading control). (**F**) Representative immunofluorescence images of U2OS WT cells 30 min after UV irradiation with 60 J/m$^2$ UV-C through a microporous filter to induce LUD. Images show accumulation of endogenous EXO1 at LUD. Coverslips were stained with antibodies against EXO1 and CPD. DNA is stained with DAPI. Scale bar 10 µm. (**G**) Immunoblot analysis of XPG-mClover U2OS DIVa cells treated with control or EXO1 siRNA. Immunoblots were stained with antibodies against EXO1 and tubulin (loading control). (**H**) Immunoblot analysis of XPG-mClover knock-in and XPG-mClover mScarlet-PCNA Knock-in U2OS DIVa cells. Immunoblot was stained with antibodies against XPG, PCNA and tubulin (loading control). (**I**) Representative images of fixed U2OS DIVa XPG-mClover mScarlet-PCNA knock-in cells. Cells with a homogenous PCNA fluorescence signal represent non-S phase cells while cells where PCNA is organized in foci represent S-phase cells. DNA was stained with DAPI. Scale bar represents 10 µM.

**Table S1.** Cell lines

| Cell line | Genotype | Reference |
|---|---|---|
| U2OS wild type | - | *ATCC* |
| U2OS | GFP-XPB | *Muniesa et al, 2024* |
| U2OS | GFP-XPB / XPA KO | *This study* |
| U2OS | XPD-mAID-mClover | *This study* |
| U2OS | XPF-TEV-mAID-mClover | *This study* |
| U2OS | XPF-TEV-mAID-mClover / XPA KO | *This study* |
| U2OS DIVa | DIVa | *Iacovoni et al, 2010* |
| U2OS DIVa | XPG-mAID-mClover | *This study* |
| U2OS DIVa | XPG(E791A)-mAID-mClover | *This study* |
| U2OS DIVa | XPG-mAID-mClover / mScarlet-PCNA | *This study* |
| U2OS DIVa | XPG-mAID-mClover / XPC KO | *This study* |
| U2OS | XPG-mAID-mClover | *This study* |
| U2OS | XPG-mAID-mClover / XPA KO | *This study* |
| U2OS | XPF KO | *Muniesa et al, 2024* |
| U2OS | XPG KO | *Muniesa et al, 2024* |

**Table S2.** Primary antibodies

| Antibody | Host | Source | Dilution | |
|---|---|---|---|---|
| | | | **WB** | **IF** |
| CPD | Ms | MBL international, TDM-2 | N.A. | 1/1000 |
| EXO1 | Rb | Abcam, ab155553 | 1/1000 | N.A. |
| EXO1 | Rb | Sigma, HPA053079 | N.A. | 1/500 |
| GFP | Rb | Abcam, Ab290 | N.A. | 1/500 |
| GTF2H1/p62 | Rb | Novus Biologicals, NBP2-38556 | 1/500 | N.A. |
| Ku70 | Ms | Santa Cruz, sc-17789 | 1/1000 | N.A. |
| PCNA | Ms | Abcam, Ab29 | 1/2000 | N.A. |
| RFC1 | Rb | Thermo, PA5-35965 | 1/2500 | N.A. |
| Tubulin | Ms | Sigma Aldrich, B512 | 1/10000 | N.A. |
| XPA | Rb | GeneTex, GTX103168 | 1/2000 | N.A. |
| XPB | Rb | Abcam, Ab190698 | 1/1000 | 1/1000 |
| XPC | Rb | Bethyl, A301-121A | 1/2000 | N.A. |
| XPD | Ms | Abcam, ab54676 | 1/1000 | N.A. |
| XPF | Ms | Santa Cruz, sc-136153 | 1/500 | N.A. |
| XPG | Rb | Bethyl, A301-484A | 1/1000 | N.A. |

**Table S3.** Secondary antibodies

| Antibody | Host | Source | Dilution | |
|---|---|---|---|---|
| | | | WB | IRDye |
| Mouse | Goat | Sigma, sab4600199 | 1/10000 | 680 |
| Mouse | Goat | Sigma, sab4600214 | 1/10000 | 770 |
| Rabbit | Goat | Sigma, sab4600200 | 1/10000 | 680 |
| Rabbit | Goat | Sigma, sab4600215 | 1/10000 | 770 |
| **Antibody** | **Host** | **Source** | **IF** | **AlexaFluor** |
| Mouse | Goat | Invitrogen, A-21424 | 1/1000 | 555 |
| Rabbit | Goat | Invitrogen, A-11034 | 1/1000 | 488 |
| Rabbit | Donkey | Invitrogen, A-31573 | 1/1000 | 647 |
